# Ecophysiology of *potentilla gracilis* douglas ex hook (rosaceae): effects of night temperature and water stress on photosynthetic gas exchange

**DOI:** 10.1101/685727

**Authors:** Madhav P. Nepal, Virginia S. Berg

## Abstract

Plants in stressful environments have evolved strategies to cope with fluctuating environmental conditions. *Potentilla gracilis*, also known as Alpine Cinquefoil, grows in alpine meadows of the Rocky Mountains (USA), and is subjected to wide ranges of temperature, light intensity and water availability on a time scale of minutes to days during the growing season. Leaves often freeze to a brittle state at night, are exposed to high radiation while still frosty, dehydrate to wilting during the following light period, and then repeat the cycle the following day. The main objective of this research was to determine the effect of night temperature on subsequent photosynthetic gas exchange in *P. gracilis*. We used a photosynthetic gas exchange system to compare assimilation and stomatal conductance from light response curves of cold-acclimated *P. gracilis* following warm and chilling nights, and for plants at different water potentials. From the light response curves, dark respiration, light compensation point, maximum assimilation, light saturation point, and inhibition of photosynthesis were determined and were compared among the same plants under varying conditions. Assimilation and stomatal conductance decreased with the fall in measurement temperature, following chilling nights, and with the severity of water stress. Low night temperature and high photon flux density during the daytime, which are very common during the growing season in the field, cause a reduction in photosynthesis of the plant. The probable underlying damage during inhibition is likely repairable indicating protection rather than damage. The cold nocturnal temperature, with its less efficient biochemical repair capabilities, may partly be responsible for the reduction in assimilation of the following day. *P. gracilis* species exhibited persistent acquired freezing tolerance; substantial photosynthetic productivity over a wide range of light intensity and temperature; and significant tolerance of, and rapid recovery from, severe drought; making a maximum use of often challenging resources.

## 1. Introduction

The most common environmental stresses in temperate and alpine habitats include those associated with cold and frost [1,2]. An exposure to low nonfreezing temperatures induces morphological and physiological changes in plants that result in the development of acquired freezing tolerance [3]. Many plants increase in freezing tolerance in response to low but non-freezing temperatures through a sophisticated reconfiguration of molecules at various levels of biological organization, a process called cold acclimation [4-8]. When wheat plants grown at normal warm temperatures are subjected to a temperature of 5 °C, they are killed, but after cold acclimation, they can survive temperatures as low as -20 °C [9]. Non-acclimated rye, for example, is killed by freezing at about −5 °C, but after a period of exposure to low nonfreezing temperature, it can survive below −30°C [10]. In freezing sensitive plants, ice formation occurs inside the cytoplasm, which kills the cell. However, ice forms extracellularly withdrawing water from inside the cell in freezing tolerant plants [11]. As the temperature goes down, ice formation accelerates. Cells of freezing tolerant plants are killed when they cannot tolerate the cellular dehydration, because of failure of the membrane [12]. Such defects include alteration in membrane lipid composition or metabolic modifications [13], changes in protein content [14], enzyme activities [15], redistribution of intracellular calcium ions [16], cellular leakage of electrolytes and amino acids, and a diversion of electron flow to alternate pathways [17]. Cold acclimated plants have a higher concentration of starch at the end of the acclimation period than non-acclimated plants. During chilling, cold acclimated plants may demonstrate an osmotic adjustment, increasing solute concentration and lowering cell sap freezing point. Osmotic adjustment included, but was not completely explained by, the accumulation of free sugars [18]. Thomas and James [19] obtained a similar result in genotypes of *Lolium perenne* that were capable of cold acclimation. At the molecular levels, exposing plants to low temperature involves hundreds of cold-induced genes causing extensive reorganization of the transcriptomes [7].

Photosynthetic gas exchange in a plant can be summarized in an equation A = R + P_gross_, where A is net CO_2_ assimilation or net photosynthesis, R is respiration and P_gross_ is gross photosynthesis [20-23]. The change in photosynthetic characteristics in response to varying temperature plays an important role in plant adaptations to different environments [24]. Assimilation is inhibited when plants are exposed to short-term low temperatures [25], which is due to accumulation of soluble saccharides and reduced orthophosphate cycling from the cytosol back to the chloroplast, where the low temperature limits the ATP synthesis required for RuBP regeneration. In addition low temperature results in the expression of dehydrins, which help to protect the plant from dehydration [26]. Suppression in assimilation in cold-stressed plants results from the combined effects of light and cold temperature. Previous studies have shown that Photosynthesis is significantly reduced in *Vitis vinifera* [27] and in jojoba (*Simmondsia chinensis*) after exposure to subfreezing temperature at night due to dark respiration [28]. The amount of dark respiration in cold acclimated plant varies from species to species. For example, the dark respiration of cold-acclimated *Larix decidua* was twice that of warm-acclimated plants at all temperatures [29].

Stomatal conductance depends on multiple factors such as rate of carboxylation, internal CO_2_ concentration, shape and size of the stomatal pores, age of the leaf, humidity, temperature, light and drought [30-32], which plays an important role in the net plant photosynthesis [20,23,33]. As plants are subjected to drier conditions, their stomatal conductance declines to maintain high water use efficiency [34]. Assimilation and stomatal conductance typically decline and internal CO_2_ concentration increases with increasing drought stress [30,34-36]. Bohl et al. [37] demonstrated the dehydration of freezing tolerant cells, and the lethal effects of excess dehydration on *Potentilla gracilis* Dougl. ex Hook var. *gracilis* (Family Rosaceae). They also showed *P. gracilis* has the ability to tolerate frost formation during the growing season responding to chilling temperatures by increasing freezing tolerance. This plant grows in a stressful habitat at an altitude of 3000m of the Rocky Mountains of the United States. A number of physical factors vary daily and seasonally in its native site. Daily temperature can range between 25 °C during the day and –5 °C at night; light levels can vary from dim (cloudy) to bright, with high UV levels. Plants may experience water stress ranging from severe drying to flooding. The plant has to face two types of dehydration in a 24 h period: nocturnal dehydration due to freezing temperatures, and diurnal dehydration due to high evaporative demand from high light intensity and high temperatures in the afternoon. The short annual growth period lasts approximately 2.5 months, during which, the plant has to complete its growth and reproduction despite being under stressful conditions. Interesting field and laboratory observations of this plant by Bohl et al. [37] showed an acquisition of freezing tolerance after exposure to chilling nights. The main objective of this research was to examine the effects of cold night temperature and water potential on photosynthetic gas exchange of *Potentila gracilis*.

## 2. Materials and Methods

### 2.1. Plant Materials and Growth Conditions

*Potentilla gracilis*, commonly known as slender cinquefoil, is native to the western and northern United States. It is a perennial subshrub with several spreading stems that grow up to 20-80 cm high, a palmately compound leaf with 7-9 oblong-elliptical leaflets, and often a flat-topped cyme of many bright yellow flowers. The environmental condition in its habitat is extremely variable during its growing period: there is a great variation in temperature between day and night, and light levels vary from minute to minute during the day. Temperatures in the growing season vary from as high as near 25 °C during the day to below 0 °C at night, rainfall from drought to flooding, and light from darkly cloudy to intensely bright [37]. Live plants were collected from the Brooklyn Lake area of the Snowy Range of the Rocky Mountains (Wyoming, USA), and grown in greenhouse at the Botanical Center of the University of Northern Iowa. Typical air temperatures were near 30 °C and 20 °C for the day and night, respectively [measured with a thermocouple meter (Model 450-ATT, OMEGA Engineering, Inc., Stamford, CT, USA)], and photon flux densities (PFDs) at midday were approximately 1300 µmol m^-2^ s^-1^ [measured with a quantum sensor and meter (LI-189, LI-COR, Lincoln, NE, USA]. Supplementary light was provided by 400W high-pressure sodium HID lamps (Voight Lighting, Philadelphia, PA, USA) to provide long days in late fall in order to maintain conditions as close to those during the growing season in the field. The plants were watered every day, once in the early afternoon in winter, but twice in the morning and again in the late afternoon in summer. The plants were fertilized with an NPK mixture (20:20:20) once a month.

### 2.2. Measuring the Effect of PFDs and Night Temperature on Photosynthetic Gas Exchange

LI-COR 6400 (Li-Cor Inc., Lincoln, NE), was used to measure net CO_2_ assimilation (A; µmol m^-2^ s^-1^) and stomatal conductance (g_s;_ mol H_2_O m^-2^ s^-1^) for photon flux densities (PFD) of 0, 25, 50, 100, 200, 400, 900, 1300, 1800, and 2400 µmol m^-2^ s^-1^, yielding light response curves that were determined in different trials for measurement temperatures of 5 °C, 10 °C, 15 °C, 20 °C and 25 °C. The measurements were taken at 20 and 25 °C on two consecutive days, then at 15, 10 and 5 °C on three following days in order. The plants of the same age were chilled for five consecutive nights at 5 °C for acquiring freezing tolerance. In the first set of experiments, three sets of four plants each were placed in greenhouse from the 6^th^ nights and after, for measuring PGE following warm nights. The night temperatures was maintained at 20 °C. In the second set of experiment, another three sets of four freeing tolerant plants were used for measuring the PGE following chilling nights (5 °C). During day, the plants were maintained in the greenhouse and at night they were kept at the chilling temperature.

### 2.3. Measuring the effect of Water Potential on PGE

On each of the three freezing tolerant plants, one healthy leaflet was used to determine water potential, and one of the remaining leaflets (7-9) of the same leaf was used to measure the PGE. The leaflet for determining water potential was sampled prior to each PGE trial. The same leaflet of a plant was used for all PGE trials. Values of A and g_s_ were measured at three water potential ranges categorized as: wet, dry and very dry each at 20°C and 15°C. Thus, three levels of plant water status were achieved by differential watering: 1) wet plants (well-watered or watered every day, with turgid leaves) that had water potential of -0.35 to –0.45 MPa, 2) dry plants (not watered for four days and often had curled leaves) with water potential of -0.85 to -1.15 MPa, and 3) very dry plants (not watered for six days (most water stressed plants with wilted leaves) with water potentials of -1.85 to – 1.95 MPa. Water potential was determined with a Scholander-type pressure chamber (Soil Moisture Equipment Corp., Santa Barbara, CA, USA).

### 2.4. Data Analysis

Dark respiration (DR), light compensation point (LCP), maximum assimilation (A_max_), light saturation point (LSP), and inhibition of photosynthesis (IP) were determined for each trial from the light response curves. DR is a value (µmol m^-2^ s^-1^), whose magnitude is equal to that of A measured in the dark. In absence of light, there is no photosynthesis, so the rate of CO_2_ release (negative A) gives the respiration rate. The value of LCP (µmol m^-2^ s^-1^) for each trial was calculated using the data from each light response curves. LCP was calculated as shown in Figure 1. The value of A_max_ (µmol m^-2^ s^-1^) for each trial was obtained by taking the highest value of A from each light response curve. LSP (µmol m^-2^ s^-1^) was the value of PFD associated with A_max_ for each curve. The data for determining IP were obtained from the two measurements of A at a PFD of 100 µmol m^-2^ s^-1^ before and after exposure to high light for each trial. IP is the reduction in A, expressed as a percentage of the first (before high light) value of A. Thus, IP = 100% H (A_before_ – A_after_)/A_before_.

**Figure 1.**
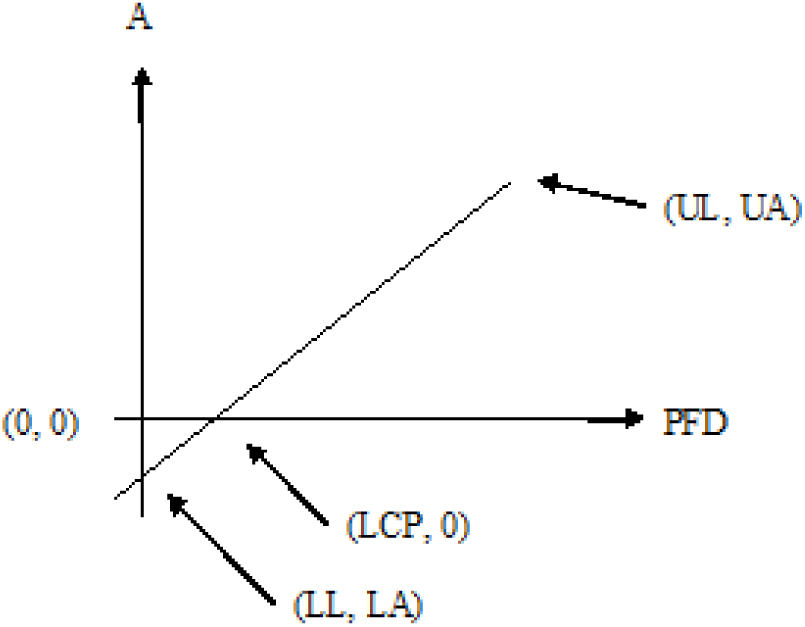
Using light response curve to calculate LCP: UL (upper light) and UA (upper A) are the values of PFD and A, respectively, for the first positive value of A measured. LL (lower light) and LA (lower A) are zero (PFD=0) and the corresponding A, respectively. General equation for a straight line is y = slope times x + b. Now, A = slope ? PFD + LA, and slope H = (UA-LA)/ (UL-LL) = ΔA/ΔL. At light compensation point, A = 0 Substituting the value of A, we get 0 = (ΔA/ΔL) H LCP + LA; -LA = (ΔA/ΔL) H LCP Thus, LCP = -LA H (ΔL/ΔA).

## 3. Results

### 3.1. Effect of Night Temperature on PGE

Following warm nights, as PFD increased, assimilation increased until the saturation point was reached, after which it remained steady or even slightly declined at higher PFDs (Figure 2). For the same PFD, assimilation was typically higher at higher temperatures. As shown in Figure 2, stomatal conductance (g_s_) increased with rising PFD. The conductance was highest at 25°C and lowest at 5°C, with intermediate values at intermediate temperatures. Following chilling nights, assimilation followed a pattern similar to that for measurements following warm nights, except that assimilation was essentially the same for the warmest two measurement temperatures. Conductance increased with rising PFD. At lower PFDs there was no clear pattern with changes in measurement temperature, but at the higher PFDs conductance was similar at lower measurement temperatures with considerably elevated for the highest two temperatures. Dark respiration (DR) following warm and chilling nights increased with the increase in measurement temperature (Figure 3, left panels). Following the two night treatments, DR levels were similar, except at 25 °C, where DR level following warm nights was considerably higher than that following chilling nights.

**Figure 2.**
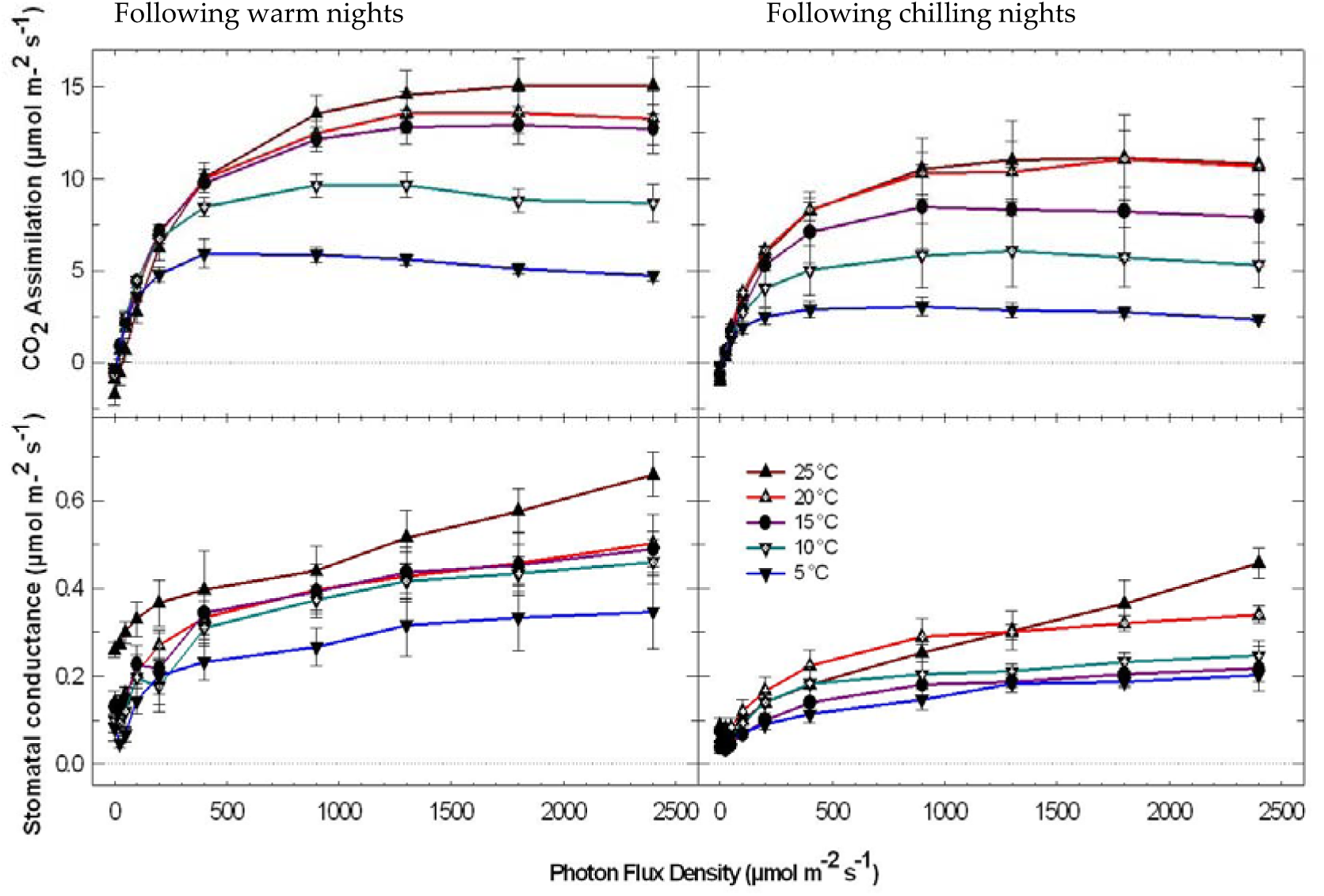
Effect of PFD on assimilation (A, upper panel left and right) and stomatal conductance (g_s_, lower panel left and right) at different measurement temperatures following warm night or chilling night as indicated. The bars represent means and standard errors.

**Figure 3.**
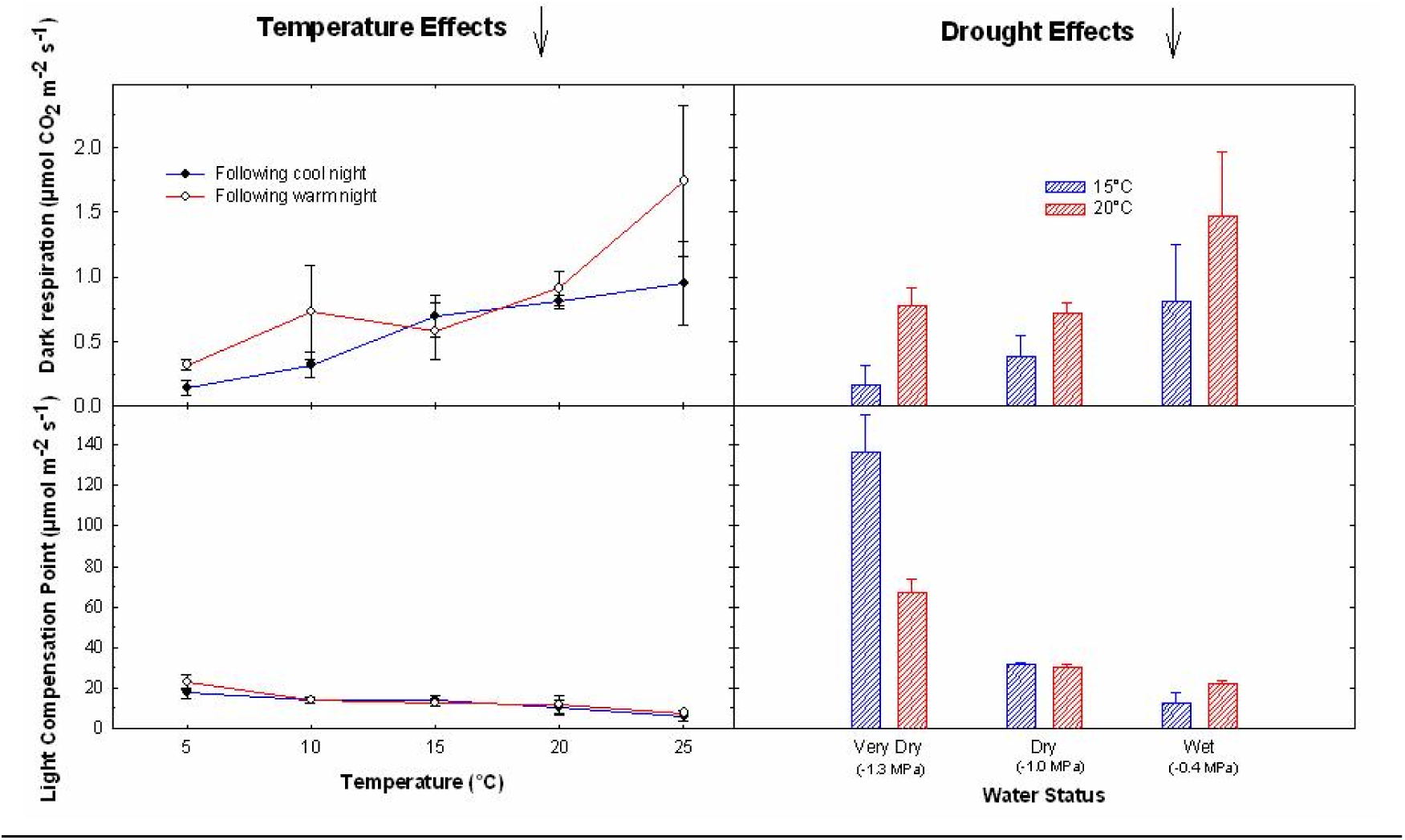
Photosynthetic Gas Exchange at 15°C and 20°C: on the left panels are the effect of night temperature on dark respiration (DR) and light compensation point (LCP), while on the right panels are the effect of water potential on DR and LCP. The bars represent means and standard errors.

Light Compensation Points (LCPs) following both warm and chilling nights decreased with increase in the measurement temperature. For all but the lowest measurement temperature, the LCP values were similar between night treatments. At 5 °C, the LCPs following warm nights were higher than those following chilling nights. Maximum Assimilation (A_max_) following warm nights increased gradually from 5 to 25 °C. Following chilling nights, A_max_ increased gradually through the range of lower measurement temperatures (5 to 20 °C), and decreased at 25 °C. At all measurement temperatures, the values of A_max_ following warm nights were higher than those following chilling nights. Light Saturation Points (LSPs) following warm nights increased progressively with increased measurement temperature. Following chilling nights, LSPs stayed approximately the same for all measurement temperatures. Inhibition of Photosynthesis (IP) declined with increased measurement temperature following warm nights. Following chilling nights, the IP was approximately constant across all the tested measurement temperatures.

### 3.2. Effect of Plant Water Potential on PGE

Assimilation at 15 °C typically increased with increasing PFD up to 900 µmol m^-2^ s^-1^ for plants at all levels of plant water status. For PFD values through 900 µmol m^-2^ s^-1^, values of A were similar for wet and dry plants but lower for very dry plants. Above 900 µmol m^-2^ s^-1^ A was highest for wet plants, intermediate for dry plants, and lowest for the very dry plants. Stomatal conductance increased as PFD increased through 400 µmol m^-2^ s^-1^, then remained nearly steady (increasing or declining slightly) at higher PFDs. Conductance was higher for the plants at wetter conditions than those under drought stress. Assimilation at 20 °C typically increased with increasing PFD up to 900 µmol m^-2^ s^-1^ for plants at the intermediate and driest conditions, above which A remained almost steady. For the wettest plants, A increased with increasing PFD through the range of PFD tested. The driest plants had positive assimilation, even below the permanent wilting point. The permanent wilting point for most plants is near -1.5 MPa (plant physiology text, but find it there). Stomatal conductance increased as PFD increased through 900 µmol m^-2^ s^-1^ for the two drier conditions, then remained nearly steady at higher PFDs. Conductance increased as PFD increased through 1300 µmol m^-2^ s^-1^ for the wet plants, and declined at higher PFDs. Values of g_s_ were highest for wet plants, intermediate for dry plants, and lowest for very dry plants.

The DR generally decreased progressively for plants under greater water stress at both 15 and 20 °C (Figure 3). The respiration was higher at 20 °C than at 15 °C for equivalent water potentials. The LCPs increased with increasing water stress at both 15 and 20 °C (Figure 3). The values of LCP were higher at 20 °C than at 15 °C for the wettest plants, similar for intermediate plants, and substantially lower for the driest plants. Maximum assimilation decreased with increased water stress at both 15 and 20 °C. The values of A_max_ were substantially higher at 20 °C than at 15 °C for the wetter two conditions, and similar for the driest condition. At 15 °C, LSP was higher for wet plants than for the two drier groups of plants (LSP derived from Figure 3). The LSPs at 20 °C decreased gradually with increasing water stress. There appeared to be only a slight increase in IP with increased water stress at 15 °C. At 20 °C, IP did not change in a regular pattern with increasing water stress. The chilled plants were equally productive if the daytime temperature was high, but if the day temperature was below 10°C, the plants did photosynthesize very low. It looks like the daytime temperature after the cold nights affect the photosynthetic yield on these plants.

## 4. Discussion

Alpine plants, including *Potentilla gracilis*, grow in harsh conditions, with wide ranging and unpredictable light intensity, plant water potential, and temperature. This plant may face dehydration in two ways everyday due to freezing nocturnal temperatures and diurnal transpiration. In the present study, this species exhibited persistent acquired freezing tolerance; substantial photosynthetic productivity over a wide range of light intensity and temperature; and significant tolerance of, and rapid recovery from, severe drought.

### 4.1. Cold Acclimation

Plant cells at temperatures that cause extracellular freezing face problems associated both with dehydration and with alteration of membrane structure and function [9,38,39]. The dehydration of cells occurs because the water potential in the intercellular spaces drops with the formation of extracellular ice. One way in which plants commonly acquire freezing tolerance is by increasing cell sap solute concentration (osmotic adjustment) by the synthesis of compatible solutes [40]. Adding solutes to the cell sap retains more water in the cell. An alternative mechanism that can be responsible for developing freezing tolerance is the synthesis of antifreeze proteins [41]. A second alternative mechanism for acquired freezing tolerance other than the accumulation of a large amount of solute is the synthesis of compounds that allow cells to tolerate dehydration and destabilizing changes in the membranes [25,42]. The synthesis of these compounds may be associated with the expression of a number of known cold induced genes [9,43].

### 4.2. Acquiring and Losing Freezing Tolerance

Freezing tolerance increased progressively as the plants were exposed to a series of consecutive chilling nights. Previously, Bohl et al. [37] predicted freezing tolerance in *P. gracilis* occurring following multiple nights at chilling temperatures. In *P. gracilis*, we evaluated the leaves damage in the plants under trial and found that leaves were no longer damaged at –6 °C after four nights at chilling temperature. Therefore, the number of days necessary for acquiring full freezing tolerance in *P. gracilis* is 4 days, which is slightly lower than in winter peas (i.e. 6 to 10 days) [44], and higher than or similar to *Arabidopsis* (which is 1 to 5 days) [40]. Acclimation to low temperature is permanent in some plants, but is partially reversible in others [45]. In *P. gracilis*, acquired freezing tolerance persisted through 45 consecutive warm nights, the end of time period examined. It is likely that it remains throughout its short growing season.

### 4.3. Photosynthetic Gas Exchange

Values of A in a typical PGE trial (at 15 °C following a warm night) increased with the increased PFD up to a point where the maximum A was reached, revealing the fact that A is light limited in that range (Figure 2). Within this range, the effect of PFD on A was a direct effect, the change in one instantly causing a parallel change in the other. This again shows that the light reaction was limiting the assimilation. As assimilation approached its maximum value, changes in A with increased PFD became less, showing the predominant limitation of the dark reaction or possible inhibition of photosynthesis due to high light. Values of g_s_ increased slowly as PFD increased stepwise (Figures 2). The maximal rate of stomatal opening was similar for each rise in PFD. The overshooting of g_s_ in response to a step change in PFD is presumably due to a drop in intercellular CO_2_ concentration (C_i_) as A increased at the higher PFD. The subsequent slight decrease in g_s_ while PFD remained constant was likely due to higher C_i_ associated with the overshoot of g_s_. Above the PFD of 1300 µmol m^-2^ s^-1^, A declined slightly, perhaps due to increased photoinhibition associated with high PFD, and to photorespiration, associated with high oxygen evolution in the light reaction. Assimilation decreased sharply at the end of the trial as PFD dropped suddenly; the corresponding value of g_s_ decreased in parallel. This must be because of the higher rate of stomatal closure than the rate of stomatal opening due to steep increases in PFD.

#### 4.3.1. Effect of Measurement Temperature on PGE following Warm Nights

When measured at 15 °C, A increased with increasing PFD, showing that A was limited by the light reaction portion of photosynthesis. At any point on the curve A consists of the sum of respiration (all forms) and gross photosynthesis (A = R + P_gross_). Dark respiration can be seen as the negative value of A at zero light. The light compensation point is the value of PFD at which the rate of respiration equals that of gross photosynthesis; thus A equals zero [31]. The quantum yield (assimilation per unit PFD, the slope of this curve) was highest between PFD levels of 0 and 25µmol m^-2^ s^-1^. Assimilation continued to increase with the increased PFD up to the first point at which A reached the maximum value. The LSP for this point was 1300 µmol m^-2^ s^-1^. Above the LSP, A was predominantly limited by the dark reaction. The quantum yield gradually decreased as the LSP was approached, showing the gradual shift from light reaction limited A to dark reaction limited A. At higher PFDs the assimilation declined slightly. This pattern is likely to be caused by a combination of photoprotection, inadequate repair of light damage to photosystem II, and photorespiration. A comparison between A at 100 µmol m^-2^ s^-1^ before and after exposure to high PFD showed that A was reduced by approximately 50%. This inhibition of photosynthesis indicates reduction in efficiency of the light reaction, likely due to conversion of xanthophylls between nonprotectant and protectant forms, and imbalance between the rates of the normal light-induced damage to photosystem II and the repair of this system [1]. The pattern of changes of g_s_ with increasing PFD followed that of A in response to PFD (Figure 4). This is expected because g_s_ and A are both directly affected by light, and because as A rises, C_i_ drops, and stomata respond to the lowered C_i_ by opening further. The result is that g_s_ adjusts to allow optimal CO_2_ for photosynthesis [1]. The value of g_s_ in *Rosa hybrida* ranged from 0.10 to 0.52 mol m^-2^ s^-1^ [46], similar to the values seen for *P. gracilis* in the present study.

**Figure 4.**
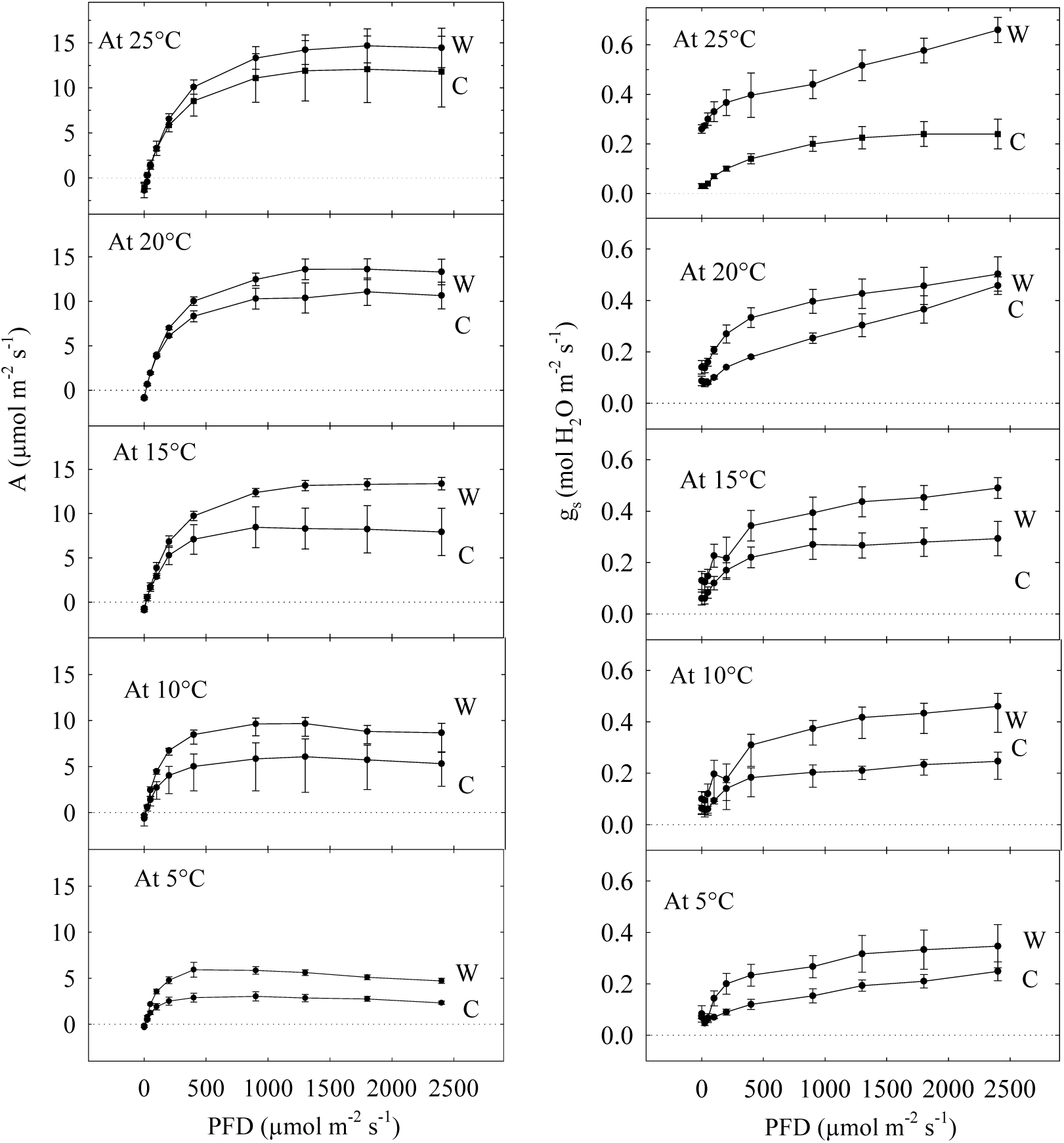
Effect of PFD and night temperature on photosynthetic assimilation (A; left) and stomatal conductance (g_s_; right) of *P. gracilis* at different measurement temperatures. W and C represent trials following warm nights and chilling nights, respectively. The bars represent means and standard errors.

At other measurement temperatures, the pattern of A with increasing PFD was generally similar to that at 15 °C. Dark respiration typically increased with the increase in measurement temperature. Respiratory enzymes become more active at higher temperatures [40]. Values of LCP decreased as measurement temperature increased. This shows that the increase in the rate of gross photosynthesis with increasing measurement temperature exceeded the corresponding increase in respiration. In the field, this can be highly significant because at higher temperatures, photosynthetic products can increase even in very low light, while at lower temperatures at the same light level, photosynthate is being consumed. For *P. gracilis* in the field, these conditions occur only momentarily on clear days, but may occur for considerable periods in morning or afternoon during cloudy or foggy days. In general, A progressively increased with increases in measurement temperature at a particular PFD. Differences between 5 and 15 °C were substantially larger than those between 15 and 25 °C. Maximum assimilation increased with increases in measurement temperature. Thus *P. gracilis* should grow considerably faster at higher temperatures in the field. Maximum assimilation for two woody *Rosa* species was 18.7 and 16.8 µmol m^-2^ s^-1^ [47], close to the values observed for *P. gracilis* at 25 °C. The PFD at which maximum A occurs (LSP) also increased with increasing measurement temperature, showing that the plants at higher temperatures were able to utilize a greater proportion of the light resource.

Inhibition of photosynthesis following exposure to high PFD declined markedly (from 61 to 38%) as measurement temperature rose. The combination of high PFD and cold temperature is expected to pose photosynthetic challenges to plants [48,49]. Photosynthetic enzyme activity is sensitive to low temperature, which affects chloroplasts directly [24]. At low temperatures the inhibition of A is due to an accumulation of soluble saccharides and reduced orthophosphate cycling from the cytosol back to the chloroplast. It limits ATP synthesis required for RuBP regeneration [24]; [26]. After chilling tomato (*Lycopersicon esculentum*) plants for a short time [50], impairment of rubisco (the enzyme that binds CO_2_) also took place, reducing A. On the other hand, as temperatures drop, the energy produced by the light reaction (a mostly physical process) cannot be fully used by the dark reaction (a mostly enzymatic process), which can lead to the damage of the photosynthetic apparatus [23]. The combination of high light and low temperature may be harmful, involving damage to photosystem II by high PFD. The repair rate of photosystem II will be lower, as the enzymes involved are less active at lower temperatures [51,52]. Light exposure at low temperature also causes an inhibition of photosystem I in barley and cucumber [53]. High temperature and high PFD have also been shown to have additive effects on photoinhibition [54], but this did not occur in the present study. The inhibition of photosynthesis in *P. gracilis* following high light may be due to the reversible conversion of xanthophylls between non-protectant (non-absorbing) and protectant (light absorbing) forms, which reduce the light getting to the chlorophyll [49,55]. This may reduce damage to the chloroplasts, at the expense of photosynthesis. The pattern of change in g_s_ with measurement temperature generally followed the pattern of change in A with measurement temperature. At temperatures from 10 to 20 °C, values overlap substantially, while those at 5 and 25 °C are considerably lower and higher, respectively, than the others. The values of g_s_ are normally higher at optimal temperatures, compared to those at colder temperatures.

#### 4.3.3. Effect of Night Temperature on PGE

Following chilling nights, the pattern of A in response to PFD was similar to that following warm nights, except that the values of A were substantially lower for corresponding PFDs and temperature levels (Figure 2). Germino and Smith [56] found that A following the warm nights was 35% higher than following cold nights for *Caltha leptosephala*, an alpine herb from same habitat as *P. gracilis*. The reduction in A after chilling nights in the present study is also in agreement with investigations of grape (*Vitis vinifera*) [27] and jojoba (*Simmondsia chinensis*) [28]. Assimilation is also inhibited in winter rye when plants are exposed to short-term (prior night) low temperatures [25] [25]. It is possible that chilling nights immediately following exposure to high PFD results in a failure to fully repair photosystem II. In contrast with the present results, Sundar and Reddy [57] found an increase in A in *Parthenium argentatum* after exposing this plant to low night temperatures.

Geiger and Servaites [24] pointed out that changes in photosynthetic characteristics in response to temperature are important in plant adaptations to different environments. For *P. gracilis*, the pattern of A in response to measurement temperature differed between trials following chilling nights and those following warm nights (Figure 2). Assimilation was nearly identical at 20 and 25 °C following chilling nights, in contrast with the pattern following warm nights, where the top three measurement temperatures had similar (but not identical) A. It is likely that the temperature optima for the limiting enzymes in the dark reaction are near 20 °C, a common temperature in the field on sunny days. In all other cases, measurement temperature had a pronounced effect, presumably because reaction rates decline at lower temperatures. Dark respiration rates following chilling nights were typically slightly lower than rates following warm nights, and also increased with measurement temperature (Figure 3). Only at 25 °C was the value of DR substantially higher (in absolute terms) for plants following warm nights. This in sharp contrast with the work of Tranquillini et al. [29], which found that dark respiration of cold-acclimated *Larix decidua* was twice that of warm-acclimated plants at all temperatures.

Values of LCP following chilling nights were almost identical to those following warm nights (Figure 3). Only at 5 °C, LCP was lower than that following warm night, a favorable response to chilling night. LCPs of sun plants range from 10-20 µmol m^-2^ s^-1^ [31], close to the values seen for *P. gracilis*. Similarly, Hopkins and Huner [23] stated that the LCPs of most plants fall between 10 and 40 µmol m^-2^ s^-1^ under favorable conditions. Others have found higher values for some woody species, such as *Rosa bracteata* and *R. rugosa* (60 and 40 µ mol m^-2^ s^-1^, respectively; [47]), and *Quercus macrocarpa* (29 to 63 µmol m^-2^ s^-1^ for shade and sun leaves, respectively [58]. These plants require high light levels to achieve positive A. Compared with values for plants measured following warm nights, A_max_ for plants following chilling nights were lower, although the increase in A_max_ with measurement temperature was parallel for the two groups, with little difference between 20 and 25 °C (Figure 2). In contrast, Hurry et al. [25] demonstrated that cold hardening increased A_max_ in winter rye (*Secale cereale*).

Values of LSP following chilling nights were essentially constant with measurement temperature, in contrast with the increasing trend with measurement temperature for plants following warm nights (Figure 3). The values coincided at 5 °C. At the whole plant level, Ballie et al. [59] reported that the light saturation point is about 1000 µmol m^-2^ s^-1^ in *Rosa hybrida*. The LSP for *P. gracilis* was nearly identical to that value following chilling nights, but following warm nights, LSP was as high as 2000 µmol m^-2^ s^-1^. Values of IP following chilling nights were similarly nearly constant, in contrast with the decreasing trend of IP with measurement temperature following warm nights (Figure 3). Values were similar between 20 and 25 °C.

In general, the pattern of g_s_ in response to PFD and measurement temperature was similar in plants following warm and chilling nights (Figures 4). For plants measured following chilling nights, g_s_ were lower than under equivalent conditions following warm nights (Figure 4). Flexas et al. [27] found similar results in grape. There was a reduction of 10 to 25 % of g_s_ in their experiment. For *P. gracilis*, this was expected, because A following chilling nights was lower than A following warm nights, and a lower g_s_ would be adequate to supply the optimum level of CO_2_. Low transpiration rates have previously been reported in chilled leaves [55]. One difference between g_s_ values for the two nocturnal temperatures is that there is significantly more overlapping of values between temperatures for plants following chilling nights.

### 4.4. Effect of Water Potentials on PGE

Assimilation at different water potentials and temperatures followed the same pattern as the previously described trials following warm and chilling nights. Assimilation was higher than at 20 °C than at 15 °C (Figure 3) for the two wetter conditions (“wet” and “dry”). However, Assimilation was essentially identical for the driest plants (“very dry”). The wet plants in these trials ranged from -.35 to -.50 MPa (−3.5 to -5.0 bars), a common value for plants in moist field sites on humid or cloudy days. The range for dry plants was -.85 to -1.10 MPa; for very dry plants (with curled leaves) water potentials ranged from -1.70 to -1.95 MPa. These values can be found for *P. gracilis* under field conditions on warm, dry days, even with adequate soil moisture. At 20 °C, A was highest for wet plants, and was not saturated at the highest PFD measured (Figure 3). Although plants typically saturate near 1000 µmol m^-2^ s^-1^ [23], some can utilize higher levels of light. For example, the LSP of *Sorghum bicolor* was 1707 and 2973 [60] for two different genotypes under wet conditions. These *P. gracilis* plants were rewatered after being subjected to very dry conditions (defined above); the high values of A on the following day show that they are able to recover quickly from severe drought stress. The dry and very dry plants were different in that A was over twice as high for the dry plants. For both groups, A saturated near 900 µmol m^-2^ s^-1^, and remained largely the same at higher PFDs. Reduction of photosynthesis may be caused by both stomatal and nonstomatal factors. Nonstomatal factors here include short-term damage to chloroplasts [36] and photosynthetic enzymes in the dark reaction that are sensitive to water potential [61]. Under drought conditions, A is controlled mostly by nonstomatal factors [30]. At 15 °C, values of A for wet and dry plants were essentially identical up to 900 µmol m^-2^ s^-1^, indicating that water potential was not limiting assimilation at these lower PFDs (Figure 3). Above this range, water potential limited assimilation; the effect was more pronounced at higher light levels. Assimilation was substantially lowered throughout the entire range of PFD for the driest plants, indicating a much greater limitation imposed by this extreme water stress. For wheat, the reduction in A due to drought stress (water potential not measured) was 65% to 80% [30]. The inhibition of photosynthesis in *P. gracilis* was approximately 43%.

At 15 °C, DR decreased as water potential declined (Figure 3), indicating the inhibition of the activity of respiratory enzymes as well those in photosynthesis. Dark respiration at 20 °C was higher for the wettest plants, and similar for the two drier groups, indicating a higher rate of metabolism of *P. gracilis* at warmer temperature and favorable plant water status. In each group, DR was higher at 20 °C than at 15 °C, in general agreement with the other trials following warm nights. The LCP increased as water potential decreased at both 15 and 20 °C, but was much more pronounced at 15 °C. The LCPs for the wet plants were higher than those observed earlier following warm nights. This may be due to a failure to recover fully in one day from severe water stress, or to the fact that the leaves used in the water stress trials were older than those in the warm night trials. Very dry plants had much higher LCPs than any other plants examined, indicating that the low water potential may have caused injury to the photosystems. The colder temperature appears to have a pronounced additional deleterious effect on the photosystems. Maximum assimilation decreased as water potential dropped. Drought stress effects on water relations of wheat Ashraf et al. [61] found that A_max_ of okra (*Hibiscus esculentus*) in dry conditions was in the range of 3.5 to 5.5 µmol m^-2^ s^-1^; in wet conditions values were in the range of 11 to 14 µmol m^-2^ s^-1^. For wheat, A_max_ was in the range of 3.9 to 4.5 µmol m^-2^ s^-1^ for dry plants, and 9 to 22 µmol m^-2^ s^-1^ for wet plants [30]. Our results are in the agreement with theirs, except that their range for water potential was such that their wet condition includes both wet and dry plants in the present study, and their dry condition was less extreme than that experience by *P. gracilis*.

Values of A_max_ were higher at 20 °C than at 15 °C for the wet plants, shifting to identical levels by the lowest water potential. The higher Amax at 20 °C for wet plants is probably due to more enzymatic activity during photosynthesis at the warmer temperature. Comparing A_max_ with the previous trials following warm nights (Figure 3), values for wet plants were identical at 20 °C, but much lower at 15 °C, equivalent to the shift from warm nights to chilling nights. This shift may be due to a slower recovery from the severe drought of the previous day. Values of LSP declined with lowered water potential, with the fastest drop from wet to dry plants at 15 °C (Figure 3). As they dried, they could not make use of available light resources. Wet plants at both temperatures had similar LSPs; the same was true for very dry plants at these temperatures. The values for wet plants were slightly higher than those for plants following the warm night trials. This means that the plants recovering from water stress were less efficient with light than those without prior water stress. The IP values were similar at all water potentials for both temperatures (Figure 3), showing that the plant was able to synthesize photoprotectants even in the presence of severe water stress. The pattern of IP was similar to that seen for the whole range of measurement temperatures following chilling nights, and the values were approximately the same. This is in contrast to the measurements following warm nights, where at the lowest temperatures; IP was substantially higher than these values.

Stomatal conductance was low, below 0.2 mol m^-2^ s^-1^, for all groups at 15 °C, and close to that value for the two drier groups at 20 °C (Figures 4). Values of g_s_ were much higher for wet plants at 20 °C. These values for *P. gracilis* were similar to those found for a wide range of plants. For okra, g_s_ under wet conditions is in the range of 0.095 to 0.150 mol m^-2^ s^-1^, while for dry plants it is 0.280 to 0.350 mol m^-2^ s^-1^ [61]. The g_s_ for two phenotypes of teak (*Tectona grandis*) varied between 0.04 to 0.32 mol m^-2^ s^-1^ for dry seedlings and between 0.35 to 0.58 µmol m^-2^ s^-1^ for wet field conditions at day temperatures from 33 to 43°C [35]. Low water potential and low temperature are characterized by lower values of g_s_ in *P. gracilus*. For wet plants at 20 °C, the high value of g_s_ is associated with a high assimilation rate, and A_max_ was the highest of all trials. The relationship between g_s_ and A is based on the fact that C_i_ controls both g_s_ and A, and in turn g_s_ and A control C_i_ [1]. As drought progresses, A decreases along with a decline in g_s_ [34]. Our results were consistent with the finding s in two species of cedar (*Cedrus atlantica* and *C. libani*) [36]. When *P. gracilis* plants were rehydrated, A and g_s_ reverted to the levels of unstressed plants. This is consistent with the findings in wheat [30] and teak [35]. This suggests that a short-term water stress brings about reversible effects. The fast recovery also suggests that the photosynthetic apparatus is highly resistant to water deficits, whereas stomatal conductance may recover more slowly. If drought continues for a long time, protein metabolism might get impaired [1,7] and plants may be unable to fully recover.

## Conclusion

*Potentilla gracilis* thrives extremely variable light, temperature and water stresses, both seasonally and day-to-day during its growing season, within which it must grow and reproduce. It survives nocturnal freezing in the growing season by acquiring persistent freezing tolerance upon exposure to chilling nights. Our results show that photosynthetic assimilation and stomatal conductance decrease with the fall in measurement temperature, following chilling nights, and with the severity of water stress. Low night temperature and high photon flux density during daytime, which are very common during the growing season in the field, cause a reduction in photosynthesis of the plant. The probable underlying damage during inhibition is likely repairable indicating protection rather than damage. The results also show that *P.gracilis* photosynthesizes significantly even at 5 °C, despite the fact that the optimum temperature is at or above 20 °C, a range common in their habitat. *P. gracilis* is able to carry out photosynthesis even when dehydrated to the point of severe wilting, and is able to recover overnight from water potentials below -1.9 MPa. Overall, *P. gracilis* species exhibited persistent acquired freezing tolerance; substantial photosynthetic productivity over a wide range of light intensity and temperature; and significant tolerance of, and rapid recovery from, severe drought; making a maximum use of often challenging resources.

## Acknowledgments

Authors acknowledge Kopila Bhattarai, Samyok Nepal, Thomas Rinehart, Jeff Church, Paul Meyerman, Larry Hilton and Mike Schultz for their assistance on the greenhouse experiments. Drs. Steve O’Kane, Jean Gerrath, Paul Whitson and Carl Thurman contributed valuable discussions on the data interpretation.

## Author Contributions

MPN performed greenhouse experiments, conducted data analysis and wrote the manuscript. VSB revised the manuscript and assisted in data analysis/interpretation.

